# Generating Minimum-Density Minimizers

**DOI:** 10.64898/2026.01.25.701585

**Authors:** Arseny Shur, Ido Tziony, Yaron Orenstein

## Abstract

Minimizers are sampling schemes which are ubiquitous in high-throughput sequencing analysis. Assuming a fixed alphabet of size *σ*, a minimizer is defined by two positive integers *k, w* and a linear order *ρ* on *k*-mers. A sequence is processed by a sliding window algorithm that chooses in each window of length *w* + *k* − 1 its minimal *k*-mer with respect to *ρ*. A key characteristic of a minimizer is its density, which is the expected frequency of chosen *k*-mers among all *k*-mers in a random infinite *σ*-ary sequence. Minimizers of smaller density are preferred as they produce smaller samples, leading to reduced runtime and memory usage in downstream applications. While the hardness of finding a minimizer of minimum density for given input parameters (*σ, k, w*) is unknown, this problem has a huge search space of (*σ*^*k*^)! and no algorithm, apart from a trivial brute-force search, is known.

In this paper, we tackle the minimum density problem for minimizers. As a first step, we formulate this problem as an ILP of size Θ(*σ*^2*k*^ + *wσ*^*w*+*k*^) and learn from experiments that an ILP solver terminates with an optimal solution only for very small *k* and *w*. We then present our main method, called OptMini, which computes an optimal minimizer in 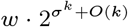 time, and is thus capable of processing large *w* values. In our experiments, OptMini works much faster than the runtime bound predicts, thanks to several additional tricks we implemented to shrink the search space without harming optimality. We use OptMini to compute minimum-density minimizers for (*σ, k*) ∈ { (2, 2), (2, 3), (2, 4), (2, 5), (2, 6), (4, 2)} over *w ∈* [2..3*σ*^*k*^], with the exception of certain *w*-ranges for *k* = 6. Finally, we derive conclusions and insights regarding the density values as a function of *w*, patterns in optimal minimizer orders, and the relation between minimum-density minimizers and minimum-size universal hitting sets.

## 1 Introduction

Sampling short substrings, termed *k*-mers, in long DNA sequences is a critical step in many bioinformatics tasks. Typically, a “window guarantee” is required: each window of *w* consecutive *k*-mers (i.e., a window of length *w* + *k* − 1) in the input sequence should be represented by at least one *k*-mer in the sample. Minimizers, introduced in [16, 18], are simple sampling schemes with the window guarantee: a linear order on *k*-mers is fixed, and in each window, the leftmost occurrence of the minimal (lowest-ranked) *k*-mer w.r.t. this order is selected. Minimizers are the most popular *k*-mer sampling schemes (a comprehensive list of bioinformatics applications using minimizers can be found in [12]), and even more advanced sampling schemes still use minimizers as an intermediate step [1, 6, 17].

Minimizers are most commonly characterized by their density, which is the expected fraction of sampled *k*-mers in an infinite sequence of i.i.d. symbols. (The related notion of *particular* density over a given *finite* sequence is not studied here.) The average density, also known as the expected density of a random minimizer, is close to 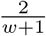 unless *w* ≫ *k* [3, 18]. Minimizers of lower density are desirable in downstream applications, as lower density leads to reduced runtime and memory usage. Consequently, many methods were designed to generate low-density minimizers. Methods like DOCKS [13], PASHA [2], and GreedyMini [4] generate orders explicitly, while methods like miniception [21], double-decycling [15], and open-closed syncmers [5] define minimizer orders via *k*-mer comparison rules, without keeping the orders themselves. However, none of the methods is guaranteed to generate nor define a minimum-density minimizer. One can only conjecture how close those methods’ results are to the minimum density, as minimum density values are known for only a few cases.

There are two trivial lower bounds on minimizers density: 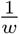 (due to window guarantee) and 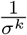 (all occurrences of the minimal *k*-mer are sampled). Both are “weakly” reachable [9]: for fixed *w*, there is an infinite sequence of minimizers with densities approaching 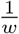 as *k* → ∞; for fixed *k*, infinitely growing *w*, and *every* order, the density approaches 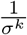. The recent KGMLT lower bound [8] is defined over forward sampling schemes, which include minimizers. It can be simplified, with a small loss in precision, to max 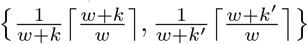, where 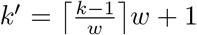 this lower bound is nearly tight for minimizers, and is even hit by minimizers for several (*k, w*) combinations [4]. However, apart from a few hits, there is still a gap in density between the lower bounds and the best known minimizers, and this gap is large for *w > k* [4, Fig. 2C,D]. Previously, no non-trivial algorithm to generate minimum-density minimizers was known, while the plain brute-force search requires iterating over *σ*^*k*^! orders, which is clearly infeasible for even small values of *k*. The related problem of finding a minimizer with the minimum particular density was considered in [20], but only for a narrow particular case of lexicographic orders.

**Figure 1.**
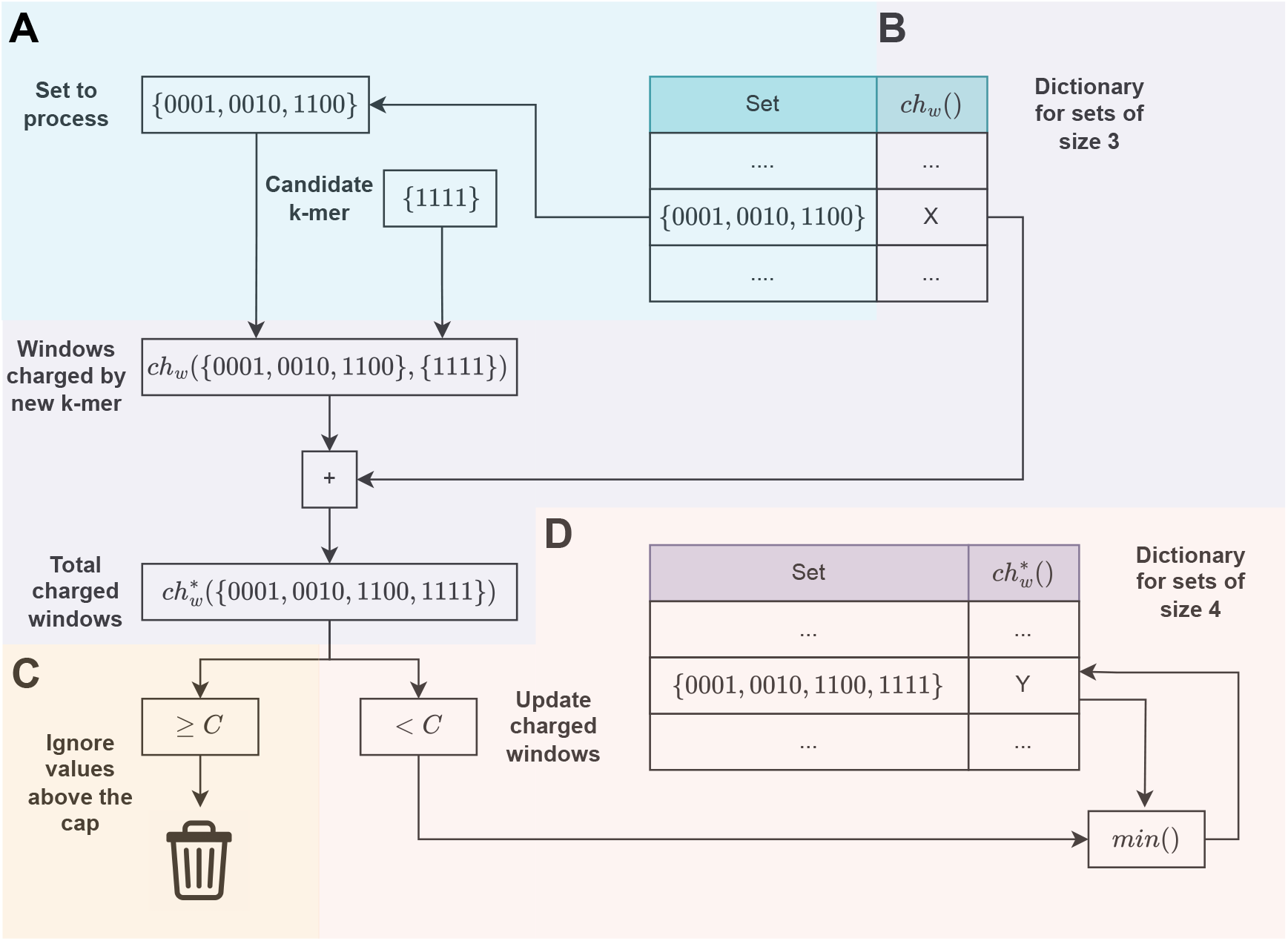
Illustration of OM-basic with the Cap trick. (A) The processed set of *k*-mers, taken from the old dictionary, and the *k*-mer added to it. (B) Windows charged due to the new *k*-mer are counted and added to the value ch_*w*_ () of the set to get an estimate of ch_*w*_ () for the extended set. (c)If the estimate hits the cap, the result is discarded as it is useless for finding an optimal order. (D)Otherwise, the estimate is used to create or update the entry in the new dictionary.

**Figure 2.**
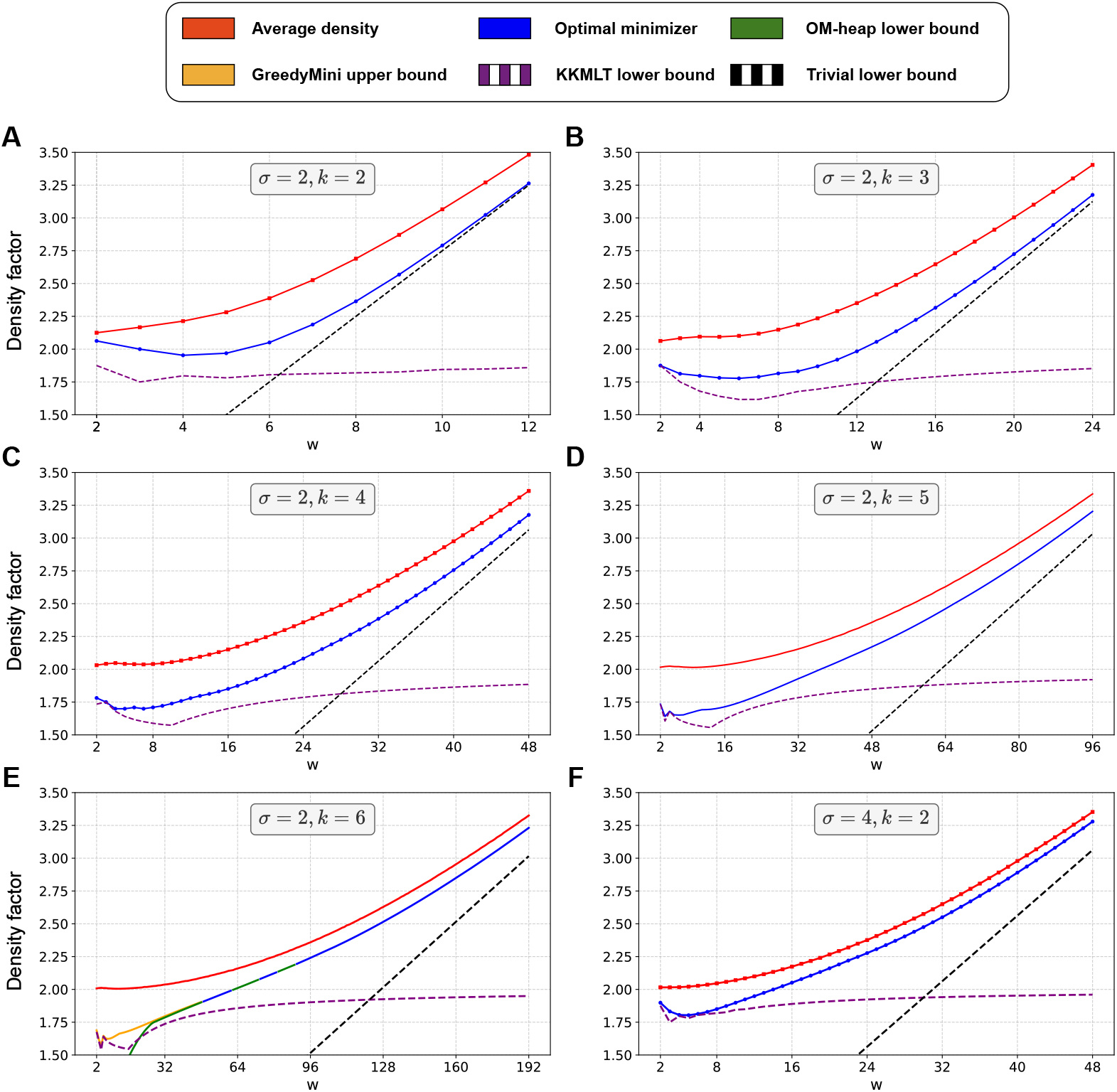
Density factors of optimal minimizers, obtained by OM-phase and OM-heap, versus lower bounds and the average density for various *k* and *σ* over *w* ∈ [2..3*σ*^*k*^]. In plot (E), whenever the optimal density is unknown, the green segments represent the lower bound obtained by OM-heap, while the yellow line is the upper bound for *w* ∈ [2..48] by the density factor of GreedyMini orders [4]. The computation of average and GreedyMini densities is specified in Appendix A.4.

In this paper, we study the problem of finding minimum-density minimizers for given parameters *k* and *w*, assuming constant *σ*. First, we formulate the problem as an ILP and show that this is insufficient to solve the problem beyond a few small cases. Our main contribution is OptMini, which finds a minimum-density minimizer in time linear in *w* and doubly exponential in *k*, assuming constant-time arithmetic operations^1^. As a by-product, OptMini can compute the average density over all minimizers in the same runtime. We describe two versions of the main algorithm of OptMini (OM-phase and OM-heap) and use their implementations to obtain minimum-density minimizer orders across wide ranges of *w* values, for the binary alphabet and *k* up to 6, and the DNA alphabet and *k* = 2. Finally, we analyze the achieved densities, observe patterns shared across optimal minimizers, and perform an additional experimental study on the relation between optimal minimizers and minimum-size universal hitting sets. See [24] for program code and tables.

## 2 Preliminaries

In what follows, Σ, *σ, k*, and *w* denote, respectively, the alphabet {0, …, *σ* − 1}, its size, the length of the sampled substrings (*k*-mers), and the number of *k*-mers in a window. We write *s*[1..*n*] for a length-*n* string, denote the length of *s* by|*s*|, and use standard definitions of substring, prefix, and suffix of a string. We write *n*-string (*n*-prefix, *n*-suffix, *n*-window) to indicate length. We use the notation [*i*..*j*] for the range of integers from *i* to *j*, and *s*[*i*..*j*] for the substring of *s* covering the range [*i*..*j*] of positions. A unary string is a string of the form *a*^*n*^, *a* ∈ Σ.

Let *ρ* be a permutation (= a linear order) of Σ^*k*^. We view *ρ* as a bijection *ρ* : [1..*σ*^*k*^] → Σ^*k*^, binding all *k*-mers to their *ρ-ranks*. The *minimizer* (*ρ, w*) is a map *f* : Σ^*w*+*k−*1^ → [1..*w*] assigning to each (*w*+*k* − 1)-window the starting position of its minimum-*ρ*-rank *k*-mer, with ties broken to the left. This map acts on strings (= sequences) over Σ, selecting one position in each window so that in a window *v* = *S*[*i*..*i*+*w*+*k* − 2] the position *i* + *f* (*v*) − 1 in *S* is selected. Minimizers are evaluated by the density of selected positions. Let *f* (*S*) denote the set of positions selected in a string *S* by a minimizer *f*. The *density of f* is the limit 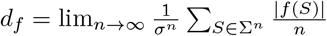 In some cases, it is convenient to “normalize” density to the *density factor D*_*f*_ = (*w* + 1)*d*_*f*_.

An *arrangement π* of Σ^*k*^ with the *domain* ∅ ≠*U* ⊆ Σ^*k*^ is an arbitrary permutation of *U*. In particular, arrangements with the domain Σ^*k*^ are exactly (linear) orders. We view *π* as a bijection *π* : [1.. |*U*|] → *U*, and apply the notion of *π*-rank to an arbitrary arrangement *π*. When *k*-mers in *π* occur in a string (e.g., in a window), the *π-minimal k*-mer is the *k*-mer of minimum *π*-rank. We also consider *π* as an ordered list of length |*π*| = |*U* |, denote the domain of *π* by *U*_*π*_, and write *π*_1_ · *π*_2_ for the arrangement obtained by concatenating arrangements *π*_1_ and *π*_2_ with disjoint domains, 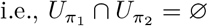.

A subset *H* ⊆ Σ^*k*^ is a *universal hitting set* (UHS) *for w* if every (*w*+*k* − 1)-window contains at least one *k*-mer from *H*. For example, { 00, 01, 11} is a UHS for *σ* = *k* = 2 and any *w >* 1. We call an arrangement *π* a *UHS order for w* if its domain *U*_*π*_ is a UHS for *w*. Then, any two orders *π· π*_1_ and *π · π*_2_ define the same minimizer for *w*, because the minimum-rank *k*-mer in every window is in *π*; we denote this minimizer by (*π, w*).

For a given minimizer *f* = (*ρ, w*), a (*w*+*k*)-window *v* (which contains two consecutive (*w*+*k* − 1)-windows) is *charged* if its minimum-*ρ*-rank *k*-mer is either its prefix or its *unique suffix* (i.e., a suffix with no other occurrence in *v*). Every string *S* contains exactly| *f* (*S*) | −1 (not necessarily distinct) charged (*w*+*k*)-windows [23, Lemma 6]. Since all possible *n*-strings have, in total, the same number of occurrences of each (*w*+*k*)-window, the density *d*_*f*_ of a minimizer equals the fraction of charged windows in Σ^*w*+*k*^ [10, 21]. For fixed *k, w*, by *window* we mean (*w*+*k*)-window. A *live* window w.r.t. a set *U* ⊂ Σ^*k*^ is a window with no *k*-mer in *U*.

For an arrangement *π* of Σ^*k*^, a window *v* is *charged by π (due to π*[*i*]) if *v* has a *k*-mer in *π*, the *π*-minimal *k*-mer of *v* is *π*[*i*] and it is a prefix (*prefix-charged*) or a unique suffix (*suffix-charged*) of *v*. The number of windows over Σ^*w*+*k*^ charged by *π* is denoted by ch_*w*_(*π*). We write ch_*w*_(*U*) = min_*π*_ {ch_*w*_(*π*) |*U*_*π*_ = *U*} and call the arrangement *π optimal for w* if ch_*w*_(*π*) = ch_*w*_(*U*_*π*_). This notion of optimality covers, in particular, optimal orders and optimal UHS orders of Σ^*k*^. Since *d*_(*ρ,w*)_ = ch_*w*_(Σ^*k*^)*/σ*^*w*+*k*^, a UHS order *ρ* is optimal for *w* if and only if the minimizer (*ρ, w*) has the minimum density among all (*σ, k, w*)-minimizers.

We use Iverson notation: [expression] evaluates to 1/0 depending on expression being true/false.

## 3 Algorithms computing optimal minimizers

### 3.1 Optimal minimizers via integer linear programming

We represent the problem of finding a minimum-density (*σ, k, w*)-minimizer with parameters as an ILP as follows. For each *k*-mer *u*, we associate a variable *x*_*u*_ ∈ [1..*σ*^*k*^] that equals the rank rank(*u*) in the constructed order. For each (*w*+*k*)-window *v*_*i*_, we associate 0-1 variables *y*_*i*_, *p*_*i*_, and *s*_*i*_ that indicate whether *v*_*i*_ is charged (respectively, prefix-charged, suffix-charged). Next, let 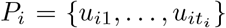 be the set of *k*-mers in *v*_*i*_, ordered by their first occurrence. We add 0-1 variables 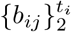that indicate whether rank(*u*_*i*1_) ≤ rank(*u*_*ij*_). In the case where *v*_*i*_ has a unique *k*-suffix (in other words, if *u*_*it*_ is a suffix of *v*_*i*_), we also add 0-1 variables 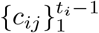 that indicate whether rank(*u*_*ij*_) *>* rank(*u*_*it*_). The total number of variables used is therefore Θ(*wσ*^*w*+*k*^). The objective is to minimize the number of charged (*w*+*k*)-windows:

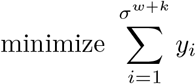

Now we define the constraints. We use boolean connectives for 0-1 values as they admit straightforward linear formulations. For each pair of variables *x*_*u*_, *x*_*v*_, we add the constraint *x*_*u*_ ≠ *x*_*v*_ as all *k*-mers have distinct ranks. We then proceed to the constraints for each (*w*+*k*)-window *v*_*i*_ separately. If *v*_*i*_ is unary, it is charged, so we just set *y*_*i*_ = 1. Otherwise, two cases are possible.

**Case 1:** the *k*-suffix of *v*_*i*_ is not unique. Then, we add *y*_*i*_ = *p*_*i*_ as *v*_*i*_ cannot be suffix-charged. By definitions of *b*_*ij*_ and prefix-charged windows, we have

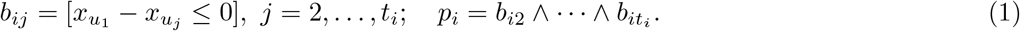

**Case 2:** the *k*-suffix of *v*_*i*_ is unique. Then, we add *y*_*i*_ = *p*_*i*_∨ *s*_*i*_. The variable *p*_*i*_ is determined by the constraints (1); in a similar way, we add the constraints defining *s*_*i*_:

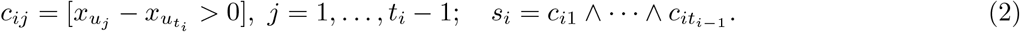

Overall, for each *v*_*i*_ we have constraints of total length *O*(*w*). Together with the pairwise non-equality constraints for the rank variables, this gives us the ILP size of Θ(*σ*^2*k*^ + *wσ*^*k*+*w*^).

### 3.2 Optimal minimizers via dynamic programming

We present OptMini, a method that performs an efficient search of a minimum-density minimizer, utilizing a combinatorial property of optimal arrangements. Suppose *σ, k*, and *w* are fixed. With every arrangement *π* of Σ^*k*^ we associate a sequence of sets 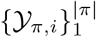 such that *Y*_*π,i*_ is the set of all (*k*+*w*)-windows charged by *π* due to *π*[*i*]. We have 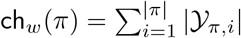. We prove the following two basic lemmas regarding arrangements.

#### Lemma 1

▸*Let π*_1_, *π*_2_ *be two arrangements with the same domain and the same last element u. Then*, 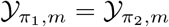, *where m* = |*π*_1_| = |*π*_2_|.

**Proof**. Let 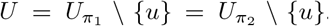. Each of the sets 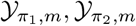 consists of all (*k*+*w*)-windows that are live w.r.t. *U* and have *u* as a prefix or a unique suffix. ◂

#### Lemma 2

▸ *If an arrangement is optimal for w, then each its prefix is optimal for w*.

**Proof**. If the lemma fails, there exists an optimal arrangement *π* = *π*_1_ (*u*) such that *π*_1_ is not optimal. Then, some arrangement *π*_2_ with the domain 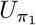 has ch_*w*_(*π*_2_) *<* ch_*w*_(*π*_1_). For *π*′ = *π*_2_ · (*u*) we have

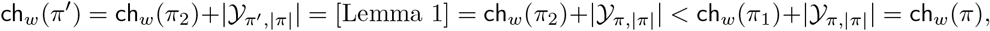

contradicting the optimality of *π*. Thus, the lemma holds. ◂

Let *π* = *π*′ · (*u*) be an arrangement of length *m*. We define ch_*w*_(*π*′, *u*) = | *Y*_*π,m*_ | to be the number of windows charged by *π* due to *u*. By Lemma 1, this number depends only on the domain 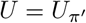 so we alternatively write ch_*w*_(*U, u*) and refer to this number as the number of windows charged due to *u after U*. For example, if *π* = (*u*_1_, …, *u*_*m*_), *U*_*i*_ = {*u*_1_, …, *u*_*i−*1_} for *i* = 1, …, *m*, then 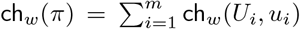. To introduce OptMini, we need the following lemma, based on the algorithms presented in [4].

#### Lemma 3

▸ *For given σ, k, w, a subset U* ⊂ Σ^*k*^, *and a k-mer u* ∈*/ U, the number* ch_*w*_(*U, u*) *can be computed in time O*(min{*σ*^*w*^, *wσ*^*k*^}).

**Proof**. Two different approaches described below result in the algorithms counting the value

▭ ch_*w*_(*U, u*) in time *O*(*σ*^*w*^) and *O*(*wσ*^*k*^), respectively. Note that the windows prefix-charged due to *u* are exactly those having the *k*-prefix *u* and all other *k*-mers outside *U* ;
▭ the windows suffix-charged due to *u* are exactly those having the *k*-suffix *u* and all other *k*-mers outside *U* ∪ {*u*}.

#### DFS Algorithm

Recall that the *trie* of a set *S* of strings is a directed rooted tree with vertices being all prefixes of strings from *S* and edges being all pairs of the form (*z, za*), where *a* ∈ Σ. Consider the trie *T* of all (*w*+*k*)-windows with the *k*-prefix *u*: it consists of the path from the root to *u* and a complete *σ*-ary tree of depth *w*, rooted at the vertex *u*. Each leaf of *T* corresponds to a window. We assume that each node *z* with |*z*| ≥ *k* is labeled by its *k*-suffix. To count windows that are prefix charged due to *u* after *U*, we set the counter ch to 0 and run a recursive DFS on the complete subtree of *T*. Visiting a vertex *v*, we check whether its *k*-suffix is in *U*. If yes, we skip the subtree of *v* (as all leaves in this subtree are not prefix-charged). Otherwise, we continue the search or, if *v* is a leaf, increment ch by 1 since *v* is prefix-charged due to *u*. By the end of the search, ch equals the number of windows prefix-charged due to *u* after *U*. For the suffix-charged windows, we consider the “dual” trie *T*′, in which all edges have the form (*z, az*), of all windows with the suffix *u*. We search it similar to searching *T*. The difference is that in suffix-charged windows *u* occurs only once. Respectively, in a vertex *v* we check whether the *k*-prefix of *v* belongs to *U* ∪ {*u*}.

The tree is not stored explicitly. Regarding strings as *σ*-ary numbers, we compute the *k*-suffixes of the children of a vertex *v* from the *k*-suffix of *v* in *O*(1) arithmetic operations (as *σ* is a constant). Then, the total number of operations in each of two DFSs is proportional to the size of the tree, i.e., to *σ*^*w*^.

#### DP Algorithm

Recall that *order-k de Bruijn graph over* Σ is a directed *σ*-regular graph, having all *σ*-ary *k*-mers as vertices and all pairs (*au, ub*), where *u* ∈ Σ^*k−*1^, *a, b* ∈ Σ, as edges. If an edge (*au, ub*) is labeled by *b*, the graph becomes a deterministic finite automaton ℬ.

We view ℬ as a *transition table* with rows indexed by *k*-mers and columns indexed by letters; the entry ℬ [*u, a*] contains the successor of *u* by the letter *a*. We take an arbitrary order *ρ* initiated by *U*. Then, we replace elements and row indices in ℬ with their *ρ*-ranks, and sort the rows by the *ρ*-rank. The resulting table is referred to asℬ _*ρ*_.

Let *n* = |*U* |+ 1. We count (*w* + *k*)-windows charged due to *u* = *ρ*[*n*] by dynamic programming. Let pref be a two-dimensional table such that pref[*r, j*] is the number of strings of length *k* + *j* having the *k*-suffix of *ρ*-rank *r*, the *k*-prefix *u*, and all other *k*-mers of *ρ*-rank at least *n*. Then, the number of (*w* + *k*)-windows prefix-charged due to *u* equals, by definition of prefix-charged, toΣ_*r≥n*_ pref[*r, w*]. In a similar way, let suff be a two-dimensional table such that suff[*r, j*] is the number of strings of length *k* + *j* having the *k*-suffix of *ρ*-rank *r* and all other *k*-mers of *ρ*-rank greater than *n*. Then, the number of (*w* + *k*)-windows suffix-charged due to *u* is suff[*n, w*].

Note that pref[*r*, 0] = [*r* = *n*], suff[*r*, 0] = [*r > n*], and the DP rules for pref and suff are almost the same: pref[*r, j* + 1] = _*ℓ*_ pref[*ℓ, j*], suff[*r, j* + 1] = _*ℓ*_ suff[*ℓ, j*], where the summation for pref (resp., suff) is over all ranks *ℓ* ≥ *n* (resp., *ℓ > n*) such that the *k*-mer of rank *r* is a successor of the *k*-mer of rank *ℓ*. Both rules can be easily computed using the transition table ℬ_*ρ*_ to propagate the counts from the *j*’th column to the (*j* + 1)th column along the edges of the de Bruijn graph. The computation requires *O*(*σ*^*k*^) time to buildℬ _*ρ*_ and *O*(*σ*^*k*^) time to compute one column of the pref and suff tables, to the total of *O*(*wσ*^*k*^). ◂

##### Theorem 4

▸ *For given σ, k, and w, a minimum-density minimizer* (*ρ, w*) *can be found in* 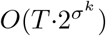 *time and* 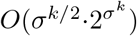 *space, where T* = min{*σ*^*w*+*k*^, *wσ*^2*k*^}.

**Proof**. Algorithm 1 (OM-basic) computes the order *ρ* by dynamic programming over subsets, proceeding in phases. At the end of the *t*’th phase, the algorithm stores 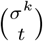 key-value pairs. The keys are all *t*-element subsets of Σ^*k*^, represented by *σ*^*k*^-bit masks; the value of the key *U* is the pair (*π*, ch_*w*_(*U*)), where *π* is an optimal arrangement with the domain *U*. Thus, at the end of the last phase the only stored value contains an optimal order *ρ*. By the definition of optimal, 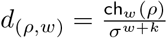 is the minimum density.

##### Algorithm 1

OM-basic: Finding an optimal order

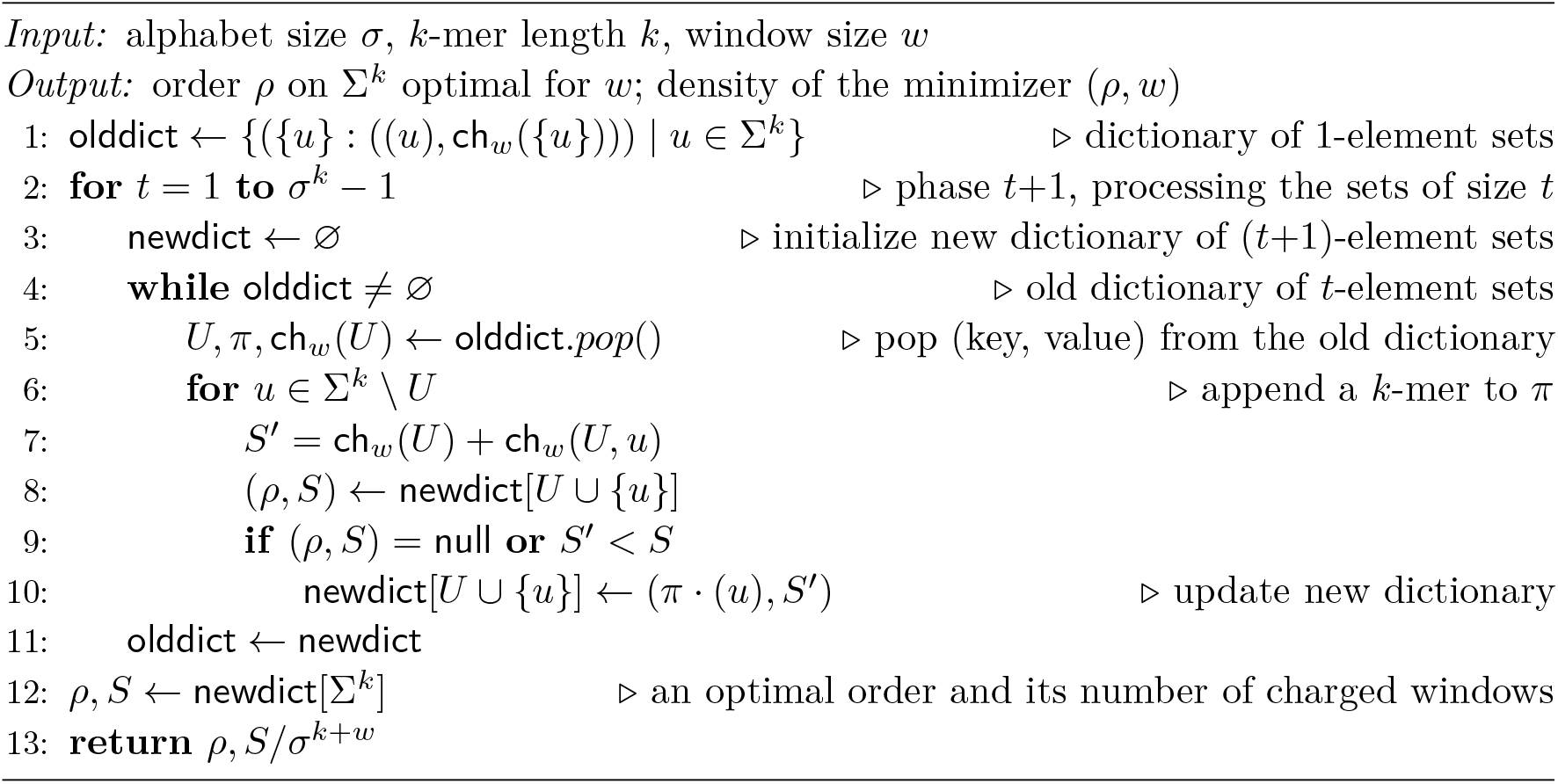

Let *U* = {*u*_1_, …, *u*_*t*_} ⊆ Σ^*k*^, *U*_*i*_ = *U\* {*u*_*i*_}, and let *π*_*i*_ be an optimal arrangement with the domain *U*_*i*_ (*i* = 1, …, *t*). We have ch_*w*_(*U*) = min{ch_*w*_(*U*_*i*_) + ch_*w*_(*U*_*i*_, *u*_*i*_) | *i* = 1, …, *t*} by Lemma 2 and the definition of ch_*w*_. If the minimum is reached on *i* = *i*^*\**^, then *π* = *π*_*i\**_ · (*u*_*i\**_) is an optimal arrangement with the domain *U*. Thus, the value for the subset *U* can be computed from the values of *t* subsets of size *t* − 1, with the runtime dominated by *t* computations of the numbers ch_*w*_(*U*_*i*_, *u*_*i*_).

In phase 1, OM-basic processes all arrangements *π* = (*u*), *u* ∈ Σ^*k*^, computing ch_*w*_({*u*}) = ch_*w*_(*π*) = |Y_*π*,1_ | and storing the value (*π*, ch_*w*_({*u*})) by the key {*u*}. During phase *t*+1, where *t* ≥ 1, it loops over all keys, which are *t*-element sets of *k*-mers (lines 4–10). Given a key *U* with the value (*π*, ch_*w*_(*U*)), it processes each *k*-mer *u ∉ U*, computing ch_*w*_(*U, u*) by means of Lemma 3 and looking up *U*′ = *U*{ *u*}in the dictionary. If no entry for *U*′ exists, it is created with the value (*π*_*u*_, ch_*w*_(*π*_*u*_)), where *π*_*u*_ = *π* · (*u*) and ch_*w*_(*π*_*u*_) = ch_*w*_(*π*) +ch_*w*_(*U, u*) = ch_*w*_(*U*) + ch_*w*_(*U, u*) by the optimality of *π*. If the entry exists (i.e., it was created earlier during this phase when processing another *t*-element subset of *U*′), its value (*τ*, ch_*w*_(*τ*)) is replaced with (*π*_*u*_, ch_*w*_(*π*_*u*_)) if ch_*w*_(*π*_*u*_) *<* ch_*w*_(*τ*). Hence, when all *t*-element subsets of *U*′ are processed, the value by the key *U*′ contains an optimal arrangement with the domain *U*′. After processing all *u ∉ U*, the key *U* is deleted. Therefore, at the end of phase *t*+1 the keys in the dictionary are exactly the (*t*+1)-element subsets of Σ^*k*^.

The dictionary dominates the space complexity of OM-basic. Since at any moment the keys are subsets of Σ^*k*^ of two consecutive sizes, the number of dictionary entries is 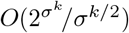 by the Stirling formula, each entry being of size *O*(*σ*^*k*^). Multiplying these numbers, we get the theorem’s space bound.

The runtime of OM-basic is dominated by the calculation of the numbers ch_*w*_(*U, u*), while the contribution of other arithmetic and dictionary operations is negligible. At most *σ*^*k*^ calculations are performed per each of 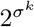 subsets. Counting *O*(min{*σ*^*w*^, *wσ*^*k*^}) per calculation (Lemma 3), we arrive at the theorem’s time bound. The theorem is proved. ◂

Having the time complexity of 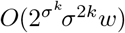 with no big constants hidden, an implementation of OM-basic is expected to terminate in a reasonable runtime for almost any *w* but for only very small *k*. For *σ* = 2, *k* ≤ 4 and for *σ* = 4, *k* = 2, where the expression under *O* evaluates to at most 224 *w*, our implementation runs within seconds, while for *σ* = 2, *k* = 5 both space and time requirements are too high. Therefore, we developed and implemented multiple tricks (Section 3.3) over OM-basic such that the resulting algorithm can cope even with the case *σ* = 2, *k* = 6, for most of the values *w* ∈ [49..3*σ*^*k*^].

### 3.3 Reducing runtime and space in the search of an optimal order

The following set of tricks dramatically reduces the runtime and space requirements for OM-basic, though their analysis does not provide improvements to the asymptotic bounds of Theorem 4. Theorem 7 guarantees correctness: the search described in Algorithm 1, enhanced with all the tricks, results in an optimal order. The combination of the first three tricks suffices to cope with the case *σ* = 2, *k* = 5, while the sixth trick makes it possible to obtain results for *σ* = 2, *k* = 6. Each trick is implemented on top of all previous ones.

#### 1. Cap trick

Having an order *ρ* on Σ^*k*^, use the value *C* = ch_*w*_(*ρ*) as a “cap”, dropping every arrangement *π* with ch_*w*_(*π*) ≥ *C* from further processing, as no order with the prefix *π* is better than *ρ* (Fig. 1). Thus, a set *U* appears in the dictionary only if some arrangement *π* with the domain *U* satisfies ch_*w*_(*π*) *< C*. The trick decreases the dictionary size and reduces the runtime accordingly. Its effect greatly increases when *ρ* is close in density to an optimal order for *w*. (In our computations, we took the best GreedyMini order [4] for each *w* ≤ 15; otherwise, we took the order *ρ* from the optimal minimizer (*ρ, w* − 1) computed beforehand.)

We also note that each of the algorithms of Lemma 3 executes different subroutines to count prefix-charged and suffix-charged windows. Hence, if the count of charged windows reaches *C* after the first subroutine, the current arrangement *π* is dropped immediately, speeding up the computation.

#### 2. UHS trick

Suppose that OM-basic, endowed with the Cap trick, processed a set *U* that contains a *k*-mer from every (*k*+*w*)-window, i.e., *U* is a UHS for *w*+1. Let *π* be an optimal arrangement with the domain *U*. Since there are no live windows w.r.t. *U*, one has ch_*w*_(*ρ*) = ch_*w*_(*U*) for every order *ρ* having *π* as a prefix^2^. This equality means that an order with ch_*w*_(*U*) charged windows is found. Hence, we reduce the remaining search by setting ch_*w*_(*U*) as a new cap (it is smaller than the previous cap, otherwise *U* would not appear in the dictionary). For more efficient use of such a “dynamic” cap, OM-basic begins processing every set *V* by comparing ch_*w*_(*V*) to the cap (which could have decreased since the moment when ch_*w*_(*V*) was computed). Processing some set *U*, the algorithm may find an optimal order; using ch_*w*_(*U*) as the upper bound may result in termination of the search before reaching the set Σ^*k*^. Accordingly, we memorize the last found UHS *U* and its dictionary value (*π*, ch_*w*_(*U*)), reporting them if the dictionary newdict is empty at the end of some phase. Note that *U* is a UHS for *w*+1 if and only if ch_*w*_(*U, u*) = 0 for all *u* ∉*U*. Since we know all the values ch_*w*_(*U, u*) after processing *U*, the trick is implemented at no cost.

If a UHS is discovered in early phases, the UHS trick reduces the number of processed sets much more efficiently. However, every UHS contains all unary *k*-mers, since it has a *k*-mer in each window (in particular, unary). Since a unary *k*-mer charges at least one window, this property postpones the moment when a UHS might be found. We circumvent this obstacle by modifying the ch_*w*_ function used in OM-basic. We define 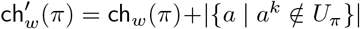 for an arrangement *π* and 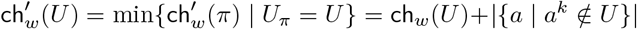 for a set *U*. Clearly, 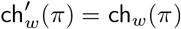 if *π* is a UHS order. Note that 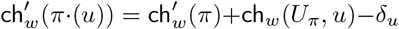, where *δ*_*u*_ = 1 if *u* is unary and *δ*_*u*_ = 0 otherwise. Hence, replacing ch_*w*_ with 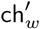 in OM-basic does not change its complexity.

We call a set *U* a *near-UHS* if all live windows w.r.t. *U* are unary. If *π* is an arrangement and *U*_*π*_ is a near-UHS, then 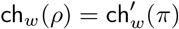 for every order *ρ* with the prefix *π*. Hence, a near-UHS *U* with an optimal arrangement *π* produces an order with 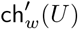 charged windows (this value is the same for any order with the prefix *π*). Therefore, 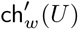 is a new cap for the remaining search. Accordingly, OM-basic stores and reports the last near-UHS it found (adding unary *k*-mers to make it a UHS) together with its value. It remains to note that *U* is a near-UHS if and only if ch_*w*_(*U, u*) − *δ*_*u*_ = 0 for all *u* ∉ *U*. Thus, after processing a set *U* we immediately know whether it is a near-UHS.

#### 3. Memory trick

By the key *U*, we store in the dictionary only the value ch_*w*_(*U*), omitting the optimal arrangement *π*. Indeed, computing ch_*w*_(*U, u*) (Lemma 3) is independent of *π*. Thus, without storing the arrangements, OM-basic still computes the optimal density. To restore an optimal order *ρ*, we use the pair 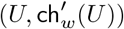 for the last found near-UHS *U*, stored due to the UHS trick. One has 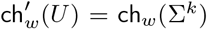 since otherwise, a near-UHS *U*^*\**^ with 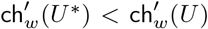 would have been found after *U*. Thus, it remains to find an optimal arrangement *π* with the domain *U*. This is done by another version of OM-basic, called OM-UHS. It uses the Cap trick, takes a set *U* as an additional input and searches for an optimal arrangement of *U* instead of Σ^*k*^. As *U* is typically significantly smaller than Σ^*k*^ (for example, in the case *σ* = 2, *k* = 5, the size of *U* over the range 3 ≤ *w* ≤ 96 is ≤ 17 out of |Σ^*k*^ | =32), the run of OM-UHS takes negligible time and space compared to the main search over Σ^*k*^.

#### 4. Sorting trick

As the dictionary olddict is never queried, we turn it into a sorted list and process sets in the order of increasing 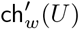 values. Paying for sorting, we gain when a near-UHS is found: the current phase can be immediately terminated as all unprocessed sets hit the new cap. Moreover, if the current set hits the cap (which may happen if the cap was decreased in the previous phase due to a near-UHS found), the current phase is terminated for the same reason. As a bonus property, among several near-UHSs of the same size, the one with the smallest 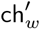 value is processed first, maximizing the decrease of the cap.

#### 5. Local Cap trick

Suppose that OM-basic employs tricks 1–4. Let *C* be the cap in the current phase (due to the Sorting trick, the change of cap terminates the phase). When a *k*-mer *u* is added to the currently processed set *U*,*U* ⋃ {*u*} is looked up in newdict *before* the computation of ch_*w*_(*U, u*). If some value 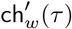) is returned (recall that due to the Memory trick, *τ* itself is not stored), then 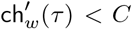, as the key *U* ⋃ {*u*} was added to newdict earlier in the current phase. Respectively, 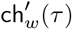 is used as the “local” cap for 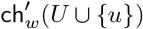 because 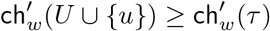 means that the value newdict[*U* ∪{ *u*}] remains unchanged. There is one important exception: if 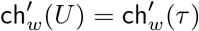, the value ch_*w*_(*U, u*) is computed in order not to miss the case where *U* is a near-UHS.

With this trick, OM-basic pays one dictionary access if 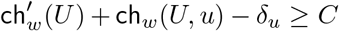 (the computed value hits the cap) for the chance to skip the whole computation or its part due to a decreased cap. Because of the Sorting trick, the first computed value for *U* ⋃{*u*} is often the smallest, which makes the local caps more efficient.

#### 6. Dead Set trick

Given a set *U* and a *k*-mer *u* ∉ *U*, we call *u wasted w*.*r*.*t. U* if *u* occurs in no live windows w.r.t. *U*. If *π* and *τ* are arbitrary arrangements with *U*_*π*_ ∩ *U*_*τ*_ = ∅, and *u* ∉ *U*_*τ*_ is wasted w.r.t. *U*_*π*_, then the arrangements *π* · *τ* and *π* · (*u*) · *τ* have exactly the same sets of charged windows and of live windows. Naturally, we would like OM-basic to process one of these arrangements rather than both. Note that if *m k*-mers satisfy the above conditions regarding *π* and *τ*, then 2^*m*^ arrangements share the sets of charged and live windows with *π · τ*, and processing just one instead of all means a great speed-up. Checking whether *u* is wasted w.r.t. *U* requires *O*(1) additional operations on top of an efficient computation of ch_*w*_(*U, u*) (Appendix A.1).

Next, we call a set *U dead* if there exists a *k*-mer *u* ∈ *U* that is wasted w.r.t. *U* \ {*u*} and satisfies 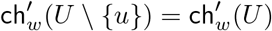 ead sets satisfy the following closure property.

##### Lemma 5

▸ *Let π and π* · *τ be optimal arrangements. If U*_*π*_ *is dead, then U*_*π·τ*_ *is dead*.

**Proof**. Let *U*_*π*_ be dead. By definition, there exists *u* ∈ *U*_*π*_ wasted w.r.t. *U*_*π*_ \ {*u*} such that 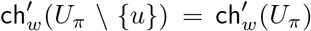 Let *π*′ be an optimal arrangement of *U*_*π*_ \ {*u*}. Then 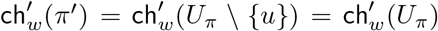, and hence *π*′ · (*u*) is optimal. Since *τ* charges the minimum possible number of windows that are live w.r.t. *U*_*π*_, the arrangement *π*′ · (*u*) · *τ* is also optimal. As *π*′ · (*u*) · *τ* and *π*′ · *τ* · (*u*) charge exactly the same windows, the latter one is also optimal. Since *u* is wasted w.r.t. *U*_*π·τ*_ \ {*u*} and 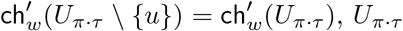 is dead by definition. ◂

A set *U* gets *labeled* if it is identified as dead while being in newdict. The search algorithm is OM-basic with tricks 1–5 and the following additional rules of processing a set *U* :

1. skip processing of *U* completely if *U* is labeled;
2. skip the computation for a *k*-mer *u* if *U* ∪{*u*} is labeled;
3. if the computation for a *k*-mer *u* is not skipped due to (ii) or the Local Cap trick, check whether *u* is wasted w.r.t. *U* ; if yes, label *U* ∪ {*u*}.

The following lemma ensures correctness of labels.

##### Lemma 6

▸ *All labeled sets are dead*.

**Proof**. Aiming at a contradiction, let *U* be a minimum-size non-dead labeled set. Let *π* = (*u*_1_, …, *u*_*n*_) be an optimal arrangement of *U* and let *π*_*i*_ = (*u*_1_, …, *u*_*i*_), *i* = 1, …, *n* − 1. The set 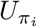 is not dead by Lemmas 2, 5 and thus not labeled by the minimality of| *U*|. Since 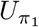 is in olddict during phase 1 and the rule (i) is not applicable to any 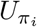, by induction each set 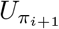 appears in newdict in phase *i* and gets the correct value 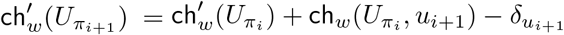. Then, *U* also gets the correct value in newdict.

Suppose that *U* is labeled while processing *U*′ = *U* \{*u*}. By (iii), *u* is wasted w.r.t. *U*′; the value computed for *U* from *U*′ is 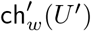. By the Local Cap trick, 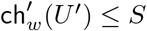 for every value *S* computed for *U* earlier. By the Sorting trick, 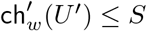 for every value *S* computed for *U* later. Since *U* at some moment gets the correct value, we have 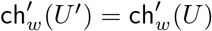 so *U* is dead by definition. The obtained contradiction proves the lemma. ◂

We refer to OM-basic enhanced with tricks 1–6 as OM-phase (Algorithm 2). Its correctness is verified by Theorem 7 below. We say that a set *U initiates* an order *ρ* if *U* = *U*_*π*_ for some prefix *π* of *ρ*.

##### Algorithm 2

OM-phase: computing minimum density and a witness UHS

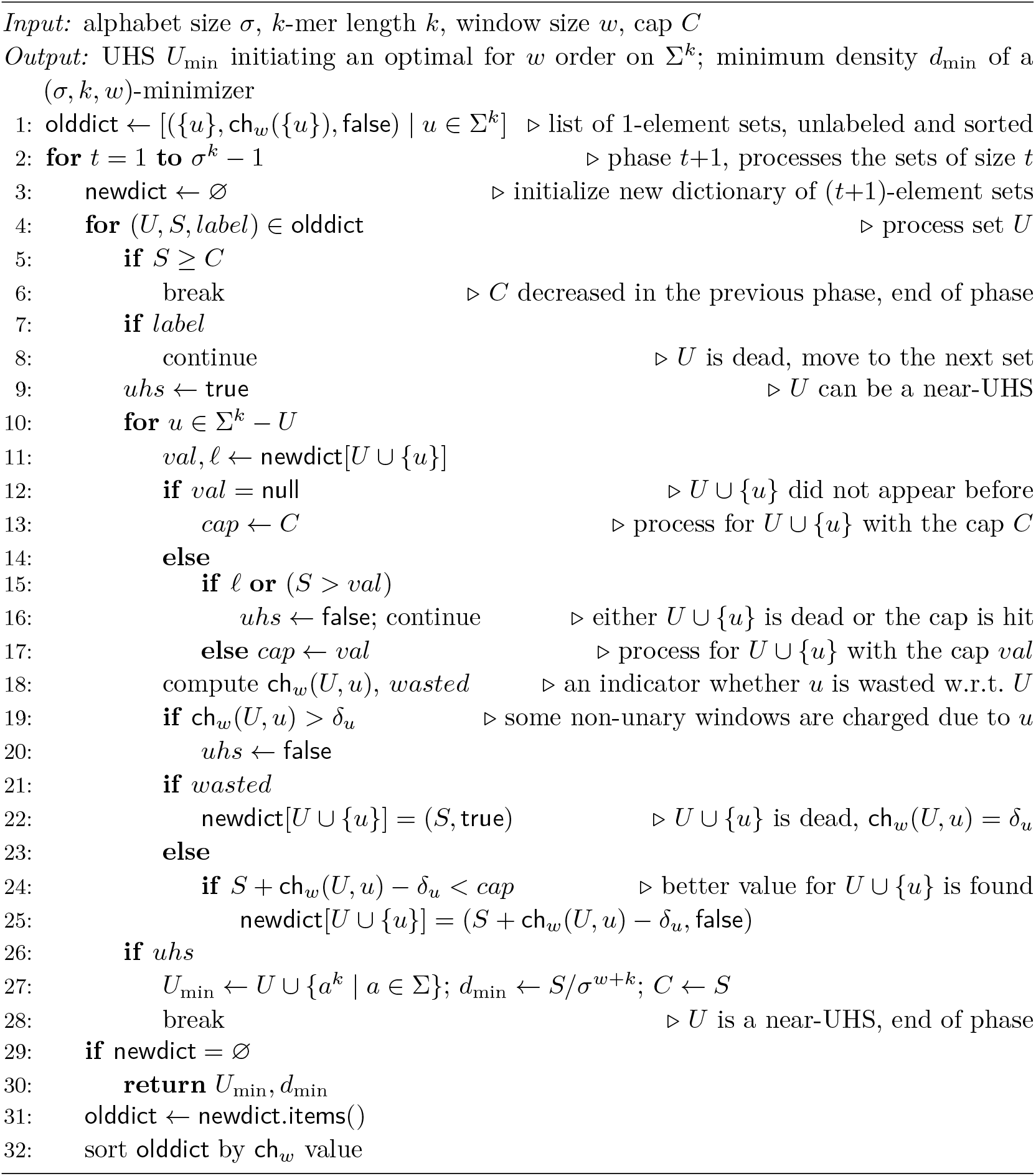

##### Theorem 7

▸ *Given σ, k, w, and a cap C >* ch_*w*_(Σ^*k*^), *OM-phase reports the minimum density among all* (*σ, k, w*)*-minimizers and a minimum-size near-UHS among all near-UHSs that initiate optimal orders for σ, k, and w*.

**Proof**. Our proof relies on the key claim showing that OM-phase correctly computes values for all sets that are both relevant to determine optimal orders and not dead.

##### Claim 8

⊳ Let *U* ⊂ Σ^*k*^, *t* = |*U* | ≥ 1, and let *C*_*t*_ be the cap during phase *t* of OM-phase. If *U* is not dead and 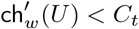 then 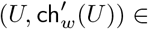 newdict at the end of phase *t*.

Proof. We prove this by induction on *t*. The base case *t* = 1 is trivial as during phase 1 newdict is populated with all pairs ({*u*}, 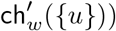 where 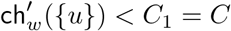.

For the step case, consider a set *U* of size *t* such that *U* is not dead and 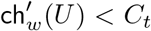. Let *π* · (*u*) be an optimal arrangement for *U*. Then,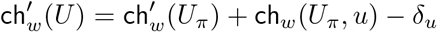. The set *U*_*π*_ is not dead by Lemma 5. As c 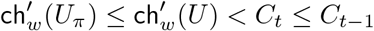, by the inductive hypothesis the pair 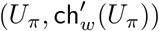 is now in olddict. As *U*_*π*_ is not labeled (Lemma 6), rule (i) of the Dead Set trick does not apply to it. Hence, *U* appears in newdict in phase *t* either during the processing of *U*_*π*_ or earlier. By Lemma 6, *U* is not labeled. So, it will get the correct value 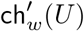 no later than during the processing of *U*_*π*_. The step case is proved.

When OM-phase processes a set *U* and finds ch_*w*_(*U, u*) = *δ*_*u*_ for every *u* ∉ *U*, it identifies *U* as a near-UHS (“discovers” *U*) and stores it with its value, later reporting the last stored near-UHS, which has smaller value than any other discovered near-UHS^3^. We say that OM-phase “misses” every near-UHS it has not discovered. To prove the theorem, it suffices to show that every near-UHS *U* satisfies at least one of the following two conditions: (a) *U* is discovered and has the value ch′_*w*_(*U*) and (b) there exists a near-UHS *U*′ such that either 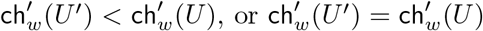 and |*U*|′ *<* |*U*|, or 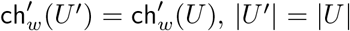, and *U*′ is discovered. (If *U* satisfies (b), missing *U* does no harm to the computation of a minimum-size optimal near-UHS order.)

Let *U* be a near-UHS, *t* = |*U*|, and let *C*_*t*_ be the cap during phase *t*. If 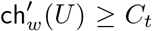, then either a near-UHS with the value *C*_*t*_ was discovered no later than phase *t* −1, or *C*_*t*_ = *C*, in which case a near-UHS with a smaller value exists. In both cases, *U* satisfies (b).

From here on, assume 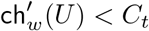 If *U* is dead, some *u* ∈ *U* is wasted w.r.t. *U*′ = *U* \{*u*} and 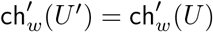 As *u* is wasted, *U*′ is a near-UHS. Hence, *U* satisfies (b).

Suppose that *U* is not dead. By Claim 8, *U* appears in newdict with the value 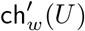. If phase *t* was terminated before reaching *U*, then some near-UHS *U*′ with |*U*′ |= |*U* |was discovered. Due to the Sorting trick, 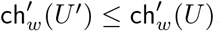, so (b) holds for *U*. Now suppose that phase *t* reached *U*. By Lemma 6, *U* is not labeled. Then, (i) is not applicable, and so *U* is processed. If ch_*w*_(*U, u*) is computed for every *u ∉ U*, then *U* is discovered and (a) holds. Hence, if *U* is missed, the computation for some *u ∉U* is skipped. If it is skipped by the Local Cap trick, then the near-UHS *U*′ = *U* ∪ {*u*} satisfies 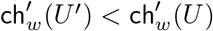, yielding the condition (b) for *U*. Otherwise, this computation is skipped by rule (ii), meaning that *U*′ is labeled and then dead (Lemma 6). Then, *U*′ = *U*^*′′*^ ∪ {*u*′}, where *u*′ is wasted w.r.t. *U*^*′′*^ (hence *U*^*′′*^ is also a near-UHS), 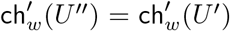 and *U*^*′′*^ is processed before *U* during the same phase (since *U*′ is already labeled when *U* is processed). We know that *U*^*′′*^ was missed, as the phase reached *U*. Hence, we repeat the above argument until after finitely many steps we arrive to a near-UHS *U*^*\**^ such that some ch_*w*_(*U*^*\**^, *u*^*\**^) was not computed due to the Local Cap trick. Then, some near-UHS has a value smaller than 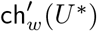. Since 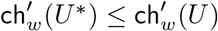 by the Sorting trick, (b) holds for *U*.

Since we have shown that every near-UHS satisfies (a) or (b), the theorem is proved. ◂

### 3.4 An alternative version: OM-heap

The OM-phase algorithm processes sets in order of their size and, within one size, in order of their 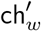 values. Along with OM-phase, we implemented an alternative version, called OM-heap, that processes sets in order of ch′_*w*_ regardless of size. OM-heap uses the UHS and Memory tricks plus a specific combination of tricks replacing the Dead Set trick. Our implementation of OM-heap employs a min-heap to store the sets to be processed (instead of olddict) and an extended version of the dictionary newdict to store the currently best values for the sets included in the heap. A detailed description and the proof of correctness of OM-heap is given in a separate section (Appendix A.2).

OM-heap never processes a set that charges more windows that an optimal order does, while OM-phase typically processes a lot of sets with values between the optimum and the cap unless the cap is set to be the optimum plus 1. On the other hand, OM-heap requires significantly more space and has to maintain a priority queue. While testing the resource-critical case *σ* = 2, *k* = 6, we used OM-heap as it obtains tighter lower bounds for those *w* where both OM-phase and OM-heap did not terminate within the allotted time.

### 3.5 A by-product: computing the average density

Given *σ, k*, and *w*, the *average density* (or *expected density of a random minimizer*) is the average of densities of all (*σ, k, w*)-minimizers. In the case *w* ≤ *k*, it can be computed in *O*(*w* log *w*) time due to a special structure of windows containing repeated *k*-mers [3], but for *w > k*, [3] gives only an *O*(*kσ*^*w*+*k*^)-time algorithm, while no algorithms that are polynomial in *k* or *w* were previously known. With an appropriate modification of OM-basic, we show the following result.

#### Theorem 9

▸ *The average density over all minimizers with parameters* (*σ, k, w*) *can be computed in* 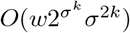 *time and O*(*σ*^*k*^) *space*.

**Proof**. Let *A* _*σ*_(*k, w*) be the average density of a minimizer with the parameters (*σ, k, w*). By definitions, we have

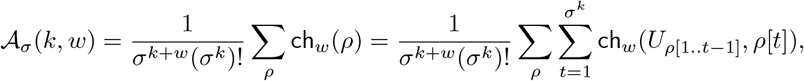

where *ρ* runs over all orders on Σ^*k*^. We swap the summation signs and write

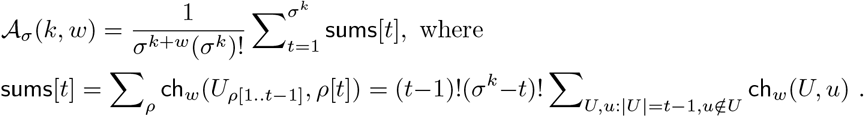

Thus, to find *A*_*σ*_(*k, w*) it suffices to compute the numbers ch_*w*_(*U, u*) for all subsets *U* ⊂ Σ^*k*^ and each *u* ∉ *U*. This computation can be performed by the dynamic-programming Algorithm 3 in *O*(*wσ*^*k*^) time per pair (*U, u*) (Lemma 3) using *O*(1) arrays of size *O*(*σ*^*k*^) each. As there is no need to store intermediate results (though one can store the array sums of size *σ*^*k*^), the described computation fits into *O*(*σ*^*k*^) space. Multiplying the time complexity of computing 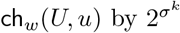 options for the set *U* and at most *σ*^*k*^ options for the *k*-mer *u*, we obtain the claimed time bound. ◂

## 4 Experimental results

### 4.1 ILP results motivate non-trivial search algorithms

We solved our ILP formulation by Gurobi [7]. We let the solver utilize all available cores on a Linux server equipped with two Intel(R) Xeon(R) Gold 6338 CPUs @ 2.00 GHz and 512 GB of RAM (64 cores total). We ran the ILP solver on values of (*σ* = 2, *w, k*), which were previously reported [8] for the wider class of *forward sampling schemes*. We limited the runtime of each run to 48 hours, seeded *x*_*i*_ variables with the ranks of *k*-mers in an order generated by GreedyMini [4], and set the solver’s lower bound to be the KGMLT bound [8]. For full numerical results, see Appendix A.3.

In all of our tests, the ILP either proved the optimality of GreedyMini orders or failed to terminate with a proven optimal solution within the time limit. In cases where the ILP terminated, the KGMLT lower bound was tight whenever *k* = 1 (mod *w*) and *k >* 1, and not tight otherwise (except for (*w, k*) = (2, 1)).

Overall, generating optimal minimizers by ILP is feasible for only a few values of *w* + *k*, in accordance with the Θ(*σ*^2*k*^ + *wσ*^*w*+*k*^) size of the ILP. This limitation calls for alternative approaches that can lift the restriction on *k* and/or *w*, and was a starting point to develop non-trivial search algorithms. The only case solved by ILP, which remained open after GreedyMini results [4] and is infeasible for OM-phase/OM-heap, is *k* = 6, *w* = 2.

### 4.2 Search algorithms reveal a part of the general picture

We computed minimum densities and optimal orders for the following minimizer parameters:

▭ *σ* = 4, *k* = 2, 2 ≤ *w* ≤ 3*σ*^*k*^ (using OM-phase)
▭ *σ* = 2, 2 ≤ *k* ≤ 5, 2 ≤ *w* ≤ 3*σ*^*k*^, except^4^ *k* = 5, *w* = 2 (using OM-phase)
▭ *σ* = 2, *k* = 6, *w* ∈ [49..61] ∪ [74..81] ∪ [90..192 = 3*σ*^*k*^] (using OM-heap)

For *σ* = 2, *k* = 6, *w* ∈ [2..48] ∪ [62..73] ∪ [82..89], OM-heap did not terminate within 24 hours on the same hardware as in Section 4.1, using one core per (*k, w*) pair. In each of these cases, the value ch_*w*_(*U*) of the last processed set gave us a lower bound for the minimum density. In order to get the optimal orders with minimum-size near-UHSs, we reproduced all minimum density results for *k* = 6 using OM-phase with the optimal cap values.

We report the density results in full in [24, Table 1] and plot them in Fig. 2 against lower bounds and the average density. The range limit *w* ≤ 3*σ*^*k*^ is not the limit of feasibility, but rather the limit of interest, since the densities of all minimizers converge to the trivial lower bound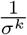as *w* approaches infinity. Below we describe the main findings from the experimental results.

#### Density behavior

Drawn in the relative scale (the density factor for the range 2 ≤ *w* ≤ 3*σ*^*k*^), all plots are similar to each other. The main density features are

□ the gap between minimum and average density factors is largest when *w* is small; for *w* ≥ *σ*^*k*^ both factors form nearly parallel curves, and the gap between them decreases as *k* or *σ* increases;
□ the rate at which minimum and average density factors approach the trivial lower bound decreases significantly as *σ* or *k* increases;
□ for minimizers, the KGMLT lower bound is not tight unless *k* = 1 mod *w* and is quite loose whenever *w > k*.

Our results for *σ* = *k* = 2 confirm the closed formula for the minimum density of a (2,2,*w*)- minimizer [19, Theorem 11].

#### Optimal orders

In the majority of studied cases, optimal orders are optimal for more than one value of *w*; e.g., one order is optimal for the range [74..96] for *σ* = 2, *k* = 5. As *k* increases, a more general pattern appears: a *family* of orders with a long prefix common to all of them “covers” a range [*w*_1_..*w*_2_]: for each *w* ∈ [*w*_1_..*w*_2_], at least one order in the family is optimal, while each of the others charges just a few more windows compared to an optimal order. For example, the range [56..66] for *σ* = 2, *k* = 5 is covered by a family of four orders that are optimal, respectively, for *w* = 56, 60, 64; *w* = 57, 61, 65; *w* = 58, 62, 66; and *w* = 59, 63; the same periodic structure is demonstrated by a family of four orders covering the range [139..147] for *σ* = 2, *k* = 6. See [24, Table 2] for details.

Similarities in the structure of optimal orders for different values of *k* can also be spotted. E.g., the optimal orders found for *σ* = 2, *k* = 3, 4, 5, 6, and *w* ∈ [2.5*σ*^*k*^..3*σ*^*k*^] all begin with the prefix (01^*k−*1^, 0^2^1^*k−*2^, …, 0^*k−*1^1).

#### Minimum density and minimum-size UHS

Previous research [9, 13, 14, 22] suggested that finding smaller UHSs leads to minimizers with lower density. Using the obtained data on optimal minimizers [24, Table 2], we tested the relation between small UHSs and minimum density. In several cases (with *σ* = 2), the orders produced by OptMini algorithms indeed have minimum-size UHSs: for *k* = 2 and any *w*; *k* = 3 and *w* = 2, 3; *k* = 4 and *w* = 3, 5; *k* = 5 and *w* = 2, 4.

However, as *w* grows, the minimum UHS size decreases monotonically to the size of a *minimum decycling set* (a UHS for *w* → ∞), while the minimum size of a near-UHS of an optimal order does not decrease (and increases on average). As a result, for large *w* there is a large gap between the minimum UHS size and the minimum size of an optimal UHS. For example, for *σ* = 2, *k* = 6 the Mykkeltveit’s minimum decycling set [11] has size 14 and is a UHS for *w >* 21. In particular, for *w* ∈ [90..192] the minimum UHS size is 14, while the minimum size of a near-UHS initiating an optimal order is 27, as was proved by OM-phase.

#### Runtime features

The actual runtime dependence on *w* is not linear; in fact, the runtime *decreases* as *w* grows (see Fig. 3 for the hardest case *k* = 6), but not monotonically: if *w* and *w*+1 are covered by the same family of orders, the runtimes are close enough; otherwise, they can differ by an order of magnitude in either direction. The fact that the cases of large *w* are easier to solve has a natural high-level explanation. If *w* is large, almost all windows contain a *k*-mer of a small rank. Hence, the density of an order depends mainly on a relatively short prefix. As a result, most of the prefixes are ruled out by the Cap trick in OM-phase or are never processed by OM-heap ; this fact greatly reduces the number of processed sets. If *w* is small, then most small-size subsets have ch _*w*_ values below the optimal number of charged windows, so both OM-heap and OM-phase process all of them.

**Figure 3.**
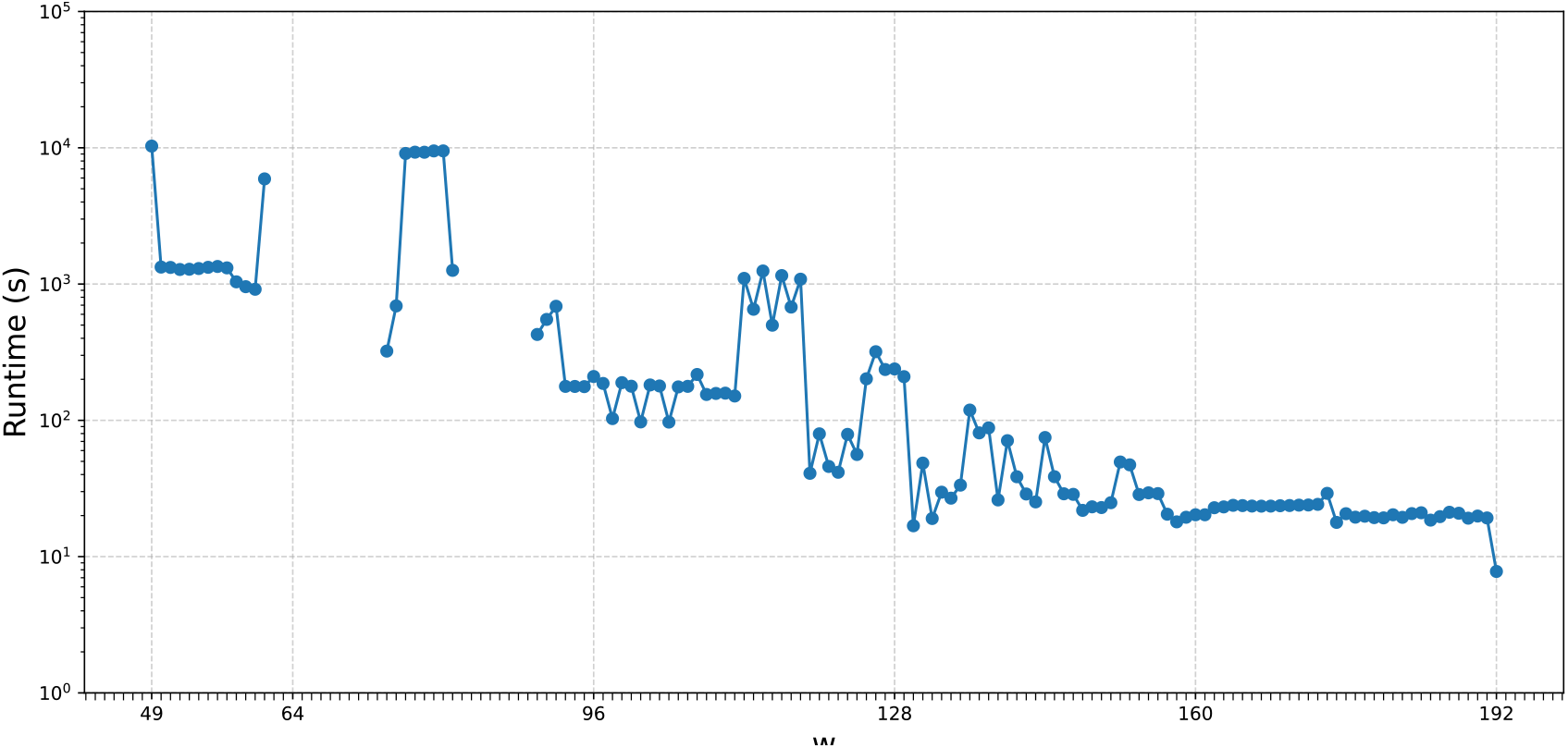
OM-heap runtime for *σ* = 2, *k* = 6 over *w* ∈ [49..3*σ*^*k*^], plotted in logarithmic scale. If a dot for some *w* value is missing, then the execution for this value did not terminate within 24 hours on the same hardware as in Section 4.1, with one instance of OM-heap per core. The time correlates well with the number of processed sets: e.g., *≈* 89,000 sets for *w* = 125 and *w* = 129, compared to *≈* 22,000 for *w* = 124 and *≈* 2,000 for *w* = 130.

## 5 Discussion

We presented OptMini, a suite of novel algorithms to generate optimal minimizers and compute their densities. The main feature of all the algorithms in OptMini is their “near independence” of the window size. While their theoretical runtime (Theorem 4) is linear in *w*, the observed runtime decreases as *w* grows. This feature allowed us to obtain a general picture of how density values and optimal orders evolve for small *σ* and *k* but over large ranges of window sizes. This is an important step in the study of minimizers: in particular, seeing the evolution of optimal orders over *w* for small *k* can result in educated guesses of low-density minimizers for practical (*k, w*) ranges.

Our study has several limitations. The runtime of the algorithms in OptMini is doubly-exponential in *k*, which severely limits the (*σ, k*) pairs they can process. By construction, OM-phase and OM-heap sort the processed sets very differently. Surprisingly, for every processed (*σ, k, w*) triple they generated UHS orders that coincide up to a few last *k*-mers. This fact suggests that the sets of optimal orders may be “homogeneous” and representable, up to symmetry, by a common prefix.

Our study raises several open questions. One of two main theoretical problems is the computational complexity of finding an optimal minimizer given a triple (*σ, k, w*). Specifically, if the runtime is polynomial in *w*, can one avoid a double exponent in *k* or prove a hardness result under standard complexity theory hypotheses? Can a better runtime bound be proved for OM-phase or OM-heap? The second main problem is proving density lower bounds. Can we prove a lower bound on the density of minimizers that is tighter than both the KGMLT and trivial bounds? Do minimizers achieve the minimum density of forward sampling schemes in infinitely many cases, for example, in all cases where *k* = *w* + 1? (It is the case for *σ* = 2 and *k* ≤ 6; see [4].)

A practically important direction for future work is studying the patterns seen in generated optimal orders and translating these patterns to practical values of *k* to obtain low-density minimizers. This work can be carried out over the binary alphabet due to the possibility of a simple translation of results to the DNA alphabet (see [4] for details), but can we gain lower density by generating orders directly to DNA alphabet, and if so, how can we do it more efficiently (in theory or practice)?

## Acknowledgements

Ido Tziony acknowledges the Bar-Ilan University Presidential Fellowship, the Planning and Budgeting Committee Data Science PhD Fellowship, and the cloud computing credit by the Israel Data Science and AI Initiative.

## A Appendix

### A.1 Faster DP computation of ch_*w*_(*U, u*) and detection of wasted *k*mers

The idea of speeding up the DP algorithm from the proof of Lemma 3 is to detect the case ch _*w*_(*U, u*) = 0 as early as possible and immediately terminate the computation. In addition, computing all the values ch _*w*_(*U, u*) in one subroutine benefits from re-using the same transition table B_*ρ*_. The pseudocode of this subroutine looks as follows.

#### Algorithm 3

Processing a set: dynamic-programming computation of ch_*w*_ (*U, u*) for all *u ∈/ U*

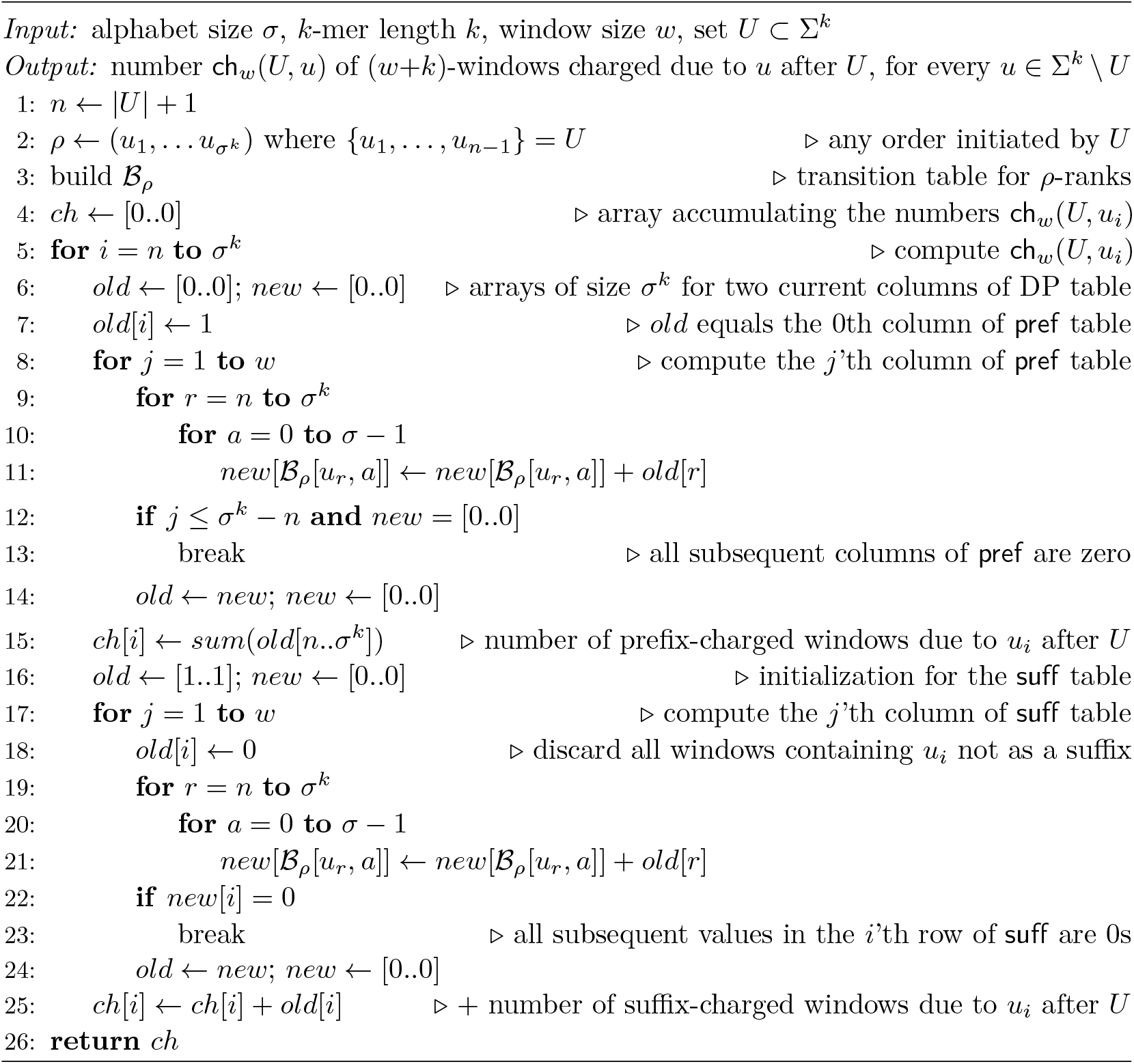

Note that all live windows w.r.t. *U* correspond to walks in the subgraph of the deBruijn graph, induced by the vertices 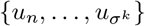; we denote this subgraph by *G*_*U*_. If *G*_*U*_ contains finitely many walks starting at *u*_*i*_, then the longest such walk is of length at most *σ*^*k*^ − *n*. Respectively, at the first *σ*^*k*^ − *n* iterations of the loop in lines 8–14 we check whether walks of length *j* from *u*_*i*_ exist; if not, then *u*_*i*_ is a prefix of no live windows, and thus we skip the rest of the computation. Similarly, if *G*_*U*_ contains finitely many walks ending at *u*_*i*_, the longest of them has length at most *σ*^*k*^ − *n*. Moreover, if no walk of length *j* ends in *u*_*i*_, then no longer walk ends in *u*_*i*_ either. Respectively, we check in line 22 whether any walk of length *j* ends in *u*_*i*_, and if not, skip the rest of the computation.

#### Detecting wasted *k*-mers

Algorithm 3 can be easily upgraded to detect wasted *k*-mers w.r.t. *U*. Recall that a *k*-mer *u* is wasted w.r.t. *U* if it occurs in no live windows. This means that, while processing *u* with Algorithm 3, we should break in both lines 13 and 23. Moreover, if we broke at the iterations *j*_1_ and *j*_2_ respectively, then the longest walks in *G*_*U*_ to/from *u* are of lengths *j*_2_ − 1 and *j*_1_ − 1 respectively. Then, *u* is wasted if and only if *j*_1_ − 1 + *j*_2_ − 1 *< w*. Note that the above check requires just *O*(1) additional operations per *k*-mer compared to Algorithm 3.

### A.2 Details of OM-heap

Algorithm OM-heap (Algorithm 4) utilizes three basic tricks. Among them, the Memory and UHS tricks are essential (though 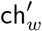 values are not used); the Cap trick reduces the size of both the heap and the dictionary, but OM-heap can also work with no cap. The other tricks described in Section 3.3 are specific for the phase-based approach, so they are replaced by additional tricks which aim to reduce the search in the heap-based approach.

The algorithm uses two main data structures:

A min-heap heap stores the sets to be processed. Each element is a triple 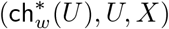, where 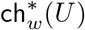 is the value computed for the set *U* at the moment of insertion into the heap. The same set can be inserted several times, each time with a strictly lower value. The set *X* is described below within the Unifying trick and is used only if *U* is a UHS.

A dictionary lowest _seen stores, by the key *U*, the lowest value 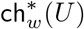 found so far. Initially, heap and lowest _seen are populated with all 1-element sets. On each iteration, OM-heap pops a minimum-valued element 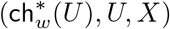 from heap and compares 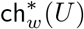 to the value stored in lowest _seen. If 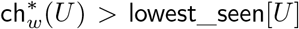, the iteration ends (*U* is discarded). Otherwise, *U* is processed. OM-heap computes the numbers ch _*w*_(*U, u*) for all *u* ∉ *U* by Algorithm 3. If *U* is a UHS (i. e., each value ch _*w*_(*U, u*) equals *δ*_*u*_), OM-heap terminates, reporting *U\X* and 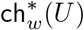. Otherwise, for each set *U′* = *U* ∪ {*u*}, the new value 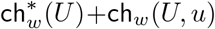 is compared to lowest _seen [*U ′*] (if the new value hits the cap, no processing is needed). If the new value is smaller or the old value does not exist, lowest _seen [*U′*] is updated, and *U′* is inserted into heap with the new value and the same *X*.

We refer to the version described above as *simple* OM-heap. It correctness follows from the order in which the heap elements are popped and Claim 10 below.

▷ Claim 10. Each processed set *U* is processed with the value ch _*w*_(*U*).

Proof. By the course of the algorithm, *U* can be inserted into heap several times, in strictly descending order of values. At the moment of last insertion, 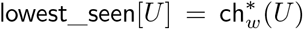 becomes the smallest of these values. In later iterations, the elements containing *U* are popped in ascending order of values; the first element is processed, while all others are discarded by comparison to 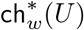. Let us prove 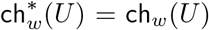 by induction on *U*. The base case |*U*| = 1 is trivial. For the step case, assume 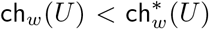 and seek a contradiction. Consider *u* ∈ *U, U′* = *U\*{*u*} such that ch _*w*_(*U*) = ch _*w*_(*U′*) + ch _*w*_(*U′, u*). As *U* is processed with the value 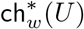, all elements with smaller values are earlier popped from heap. In particular, on some iteration *U′* is processed with its smallest value 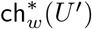, which equals ch _*w*_(*U′*) by the induction hypothesis. Then, on this iteration *U* is inserted into heap with the value, contradicting the definition of 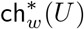 as the last value with which *U* was inserted. Hence, 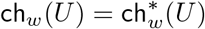, and the step case is proved. ◁

To obtain the OM-heap algorithm, we enhance the above simple version with the following two tricks reducing the search space.

#### Unifying trick

We call a *k*-mer *u* ∉ *U free w*.*r*.*t. U*, if ch _*w*_(*U, u*) = *δ*_*u*_. In particular, wasted *k*-mers are free w.r.t. the same set.

The trick is aimed at merging sets that share the same set of live windows, inflating each processed set with free *k*-mers. To balance this effect, with every set *U* we store in heap its subset *X* such that *X* is the set of wasted *k*-mers w.r.t. *U\X*. It is used to reduce the size of the reported UHS, as smaller UHS means less work to OM-UHS to recover the UHS order. Consider an iteration processing the element (ch _*w*_(*U*), *U, X*). When computing the numbers ch _*w*_(*U, u*), we keep track of the sets of free and wasted *k*-mers w.r.t. *U*. If the set *F* of free *k*-mers is empty, we proceed normally with the sets of the form *U* ∪ {*u*}. Otherwise, instead of the normal procedure we try to add *U* ∪ *F* to heap and then end the iteration. Note that the new value for *U* ∪ *F* is ch _*w*_(*U*). If it is smaller than the old value, we insert (ch _*w*_(*U*), *U* ∪ *F, X′*) into heap, where *X′* is the extension of *X* with all wasted *k*-mers found^5^.

Let us illustrate how the trick merges sets. Let *L* be the set of live windows w.r.t. some set *U* ⊂ Σ^*k*^. Let *A* be the set of all *k*-mers that are prefixes and/or unique suffixes of the non-unary windows from *L*. Then, Σ^*k*^ is partitioned into disjoint sets *U, A*, and *F*, where *F* is the set of free *k*-mers w.r.t. *U*. If *F* ≠ ∅, *U* is not processed normally but replaced in heap with *U* ∪ *F* = Σ^*k*^ \*A*. This resulting set depends only on *L*; hence the processing of any set *U′* with the same set *L* of live windows results in the same set Σ^*k*^ \*A*.

The correctness of the Unifying trick is justified by the following

##### Claim 11

▷ The Unifying trick does not change the value reported by OM-heap.

Proof. Suppose that the simple OM-heap returned the UHS *U* and thus ch _*w*_(*U*) = ch _*w*_(Σ^*k*^) is the optimal value. Let (*u*_1_, …, *u*_*t*_) be the optimal arrangement of *U* such that for each *i* = 2, …, *t*, the set *U*_*i*_ = {*u*_1_, …, *u*_*i*_} was inserted into heap with the value ch _*w*_(*U*_*i*_) when processing *U*_*i−*1_. Below we prove by induction on *i* that with the Unifying trick, for each *i* = 1, …, *t*, OM-heap processes some set *Ū*_*i*_ ⊇ *U*_*i*_ such that 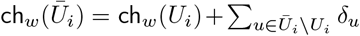. Since *U* = *U*_*t*_ is a UHS, *Ū*_*t*_ is a UHS and ch _*w*_(*Ū*_*t*_) = ch _*w*_(*U*). Hence, the algorithm terminates, at the latest, at the iteration processing *Ū*_*t*_, and thus reports the optimal value ch _*w*_(*U*).

The base case of the induction is trivial as OM-heap processes all 1-element sets. For the step case, consider *U*_*i*_. If *u*_*i*_ is free w.r.t. *U*_*i−*1_, then it is free w.r.t. a larger set *Ū*_*i−*1_ (or *u*_*i*_ ∈ *Ū*_*i*_*−*_1_, in which case we just define *Ū*_*i*_ = *Ū*_*i*_*−*_1_). By the hypothesis, *Ū*_*i*_*−*_1_ is processed, and, as having free *k*-mers, it is extended by all free *k*-mers, including *u*_*i*_. We take the result of this extension as *Ū*_*i*_: it contains *U*_*i*_ and satisfies, by the definition of free *k*-mer,

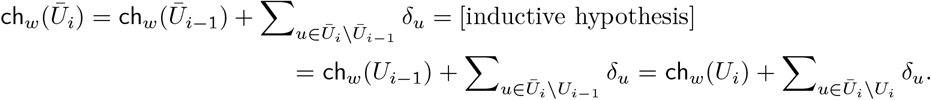

Now assume *u*_*i*_ is not free. Then, ch _*w*_(*U*_*i−*1_, *u*_*i*_) ≥ ch _*w*_(*Ū*_*i−*1_, *u*_*i*_), as all the *k*-mers live w.r.t. *Ū*_*i−*1_ are live w.r.t. *U*_*i−*1_. In case of strict inequality, we would get an impossible condition ch _*w*_(*Ū*_*i−*1_ ∪ {*u*_*i*_, …, *u*_*t*_}) *<* ch _*w*_(*U*). Hence, ch _*w*_(*U*_*i−*1_, *u*_*i*_) = ch _*w*_(*Ū*_*i−*1_, *u*_*i*_); in particular, *u*_*i*_ is not free w.r.t. *Ū*_*i−*1_. When *Ū*_*i−*1_ is processed, if no *k*-mer is free w.r.t. it, then *Ū*_*i*_ = *Ū*_*i−*1_ ∪ {*u*_*i*_} is inserted into heap with the correct value

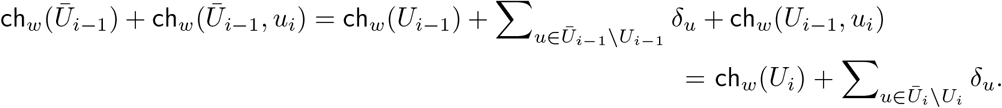

If free *k*-mers w.r.t. *Ū*_*i−*1_ exist, *Ū*_*i−*1_ is extended to a larger set with the same ch _*w*_ value. We repeat the above argument to this larger set; after finitely many repetitions we will arrive at some set *U*^*\**^ such that

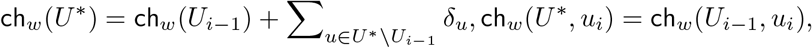

and no *k*-mers are free for *U*^*\**^. Then, *Ū*_*i*_ = *U*^*\**^ ∪ {*u*} is inserted into heap with the value 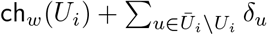 when processing *U*^*\**^. The step case is proved. ◁

#### Bad *k*-mers trick

We call a *k*-mer *u* ∉ *U bad w*.*r*.*t. U* if its in-degree or out-degree in the order-*k* de Bruijn graph induced by Σ^*k \*^*U* is 0 (Figure A4). In other words, a bad *k*-mer charges all windows in which it appears. Given that we aim at minimizing the number of charged windows, any bad *k*-mer is an unattractive choice. Accordingly, OM-heap uses the following rule: processing a set *U*, skip adding bad *k*-mers to *U* unless all *k*-mers are bad. This rule is justified by the following claim.

**Figure A4.**
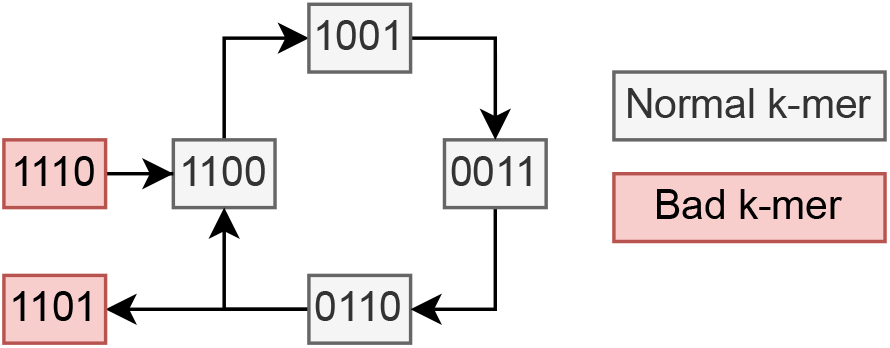
Illustration of bad *k*-mers in an order-4 de Bruijn graph induced by {0, 1}^4^ \*U* for some *U*. Each path of length *w* corresponds to one live window. Any path hitting a bad *k*-mer either begins or ends with it.

##### Claim 12

▷ Let *π* and *τ* be arbitrary arrangements with disjoint domains, and let *u* ∉ *U*_*τ*_ be bad w.r.t. *U*_*π*_. Then, ch _*w*_(*π* · *τ* · (*u*)) ≤ ch _*w*_(*π* · (*u*) · *τ*).

Proof. Let *ρ*_1_ = *π* · *τ* · (*u*), *ρ*_2_ = *π* · (*u*) · *τ*. Partition the live windows w.r.t. *U*_*π*_ into two sets: the windows containing / not containing *u*. In the non-containing set, *ρ*_1_ and *ρ*_2_ charge the same windows, while *ρ*_2_ charges all windows of the containing set. The claim is now immediate. ◁

#### Pseudocode and final remarks

Below is the pseudocode of OM-heap (Algorithm 4). For simplicity, we removed the Cap trick and the handling of unary free *k*-mers. We recall that, due to the Memory trick (Section 3.3), OM-heap only finds a UHS *U* without its optimal arrangement, and relies on OM-UHS for reconstructing the order.

The main advantage of OM-heap over OM-phase is that the former algorithm never processes a set *U* with ch _*w*_(*U*) *>* ch _*w*_(Σ^*k*^) (however, it stores such sets in both the heap and the dictionary, because ch _*w*_(Σ^*k*^) is not known until the algorithm terminates). This advantage results, in particular, in better lower bounds for the minimum density in the cases where both OM-heap and OM-phase do not terminate in reasonable time. On the other hand, OM-heap uses more memory than OM-phase and spends additional time to maintain the heap.

##### Algorithm 4

OM-heap: Heap version for finding (a UHS for) an optimal order

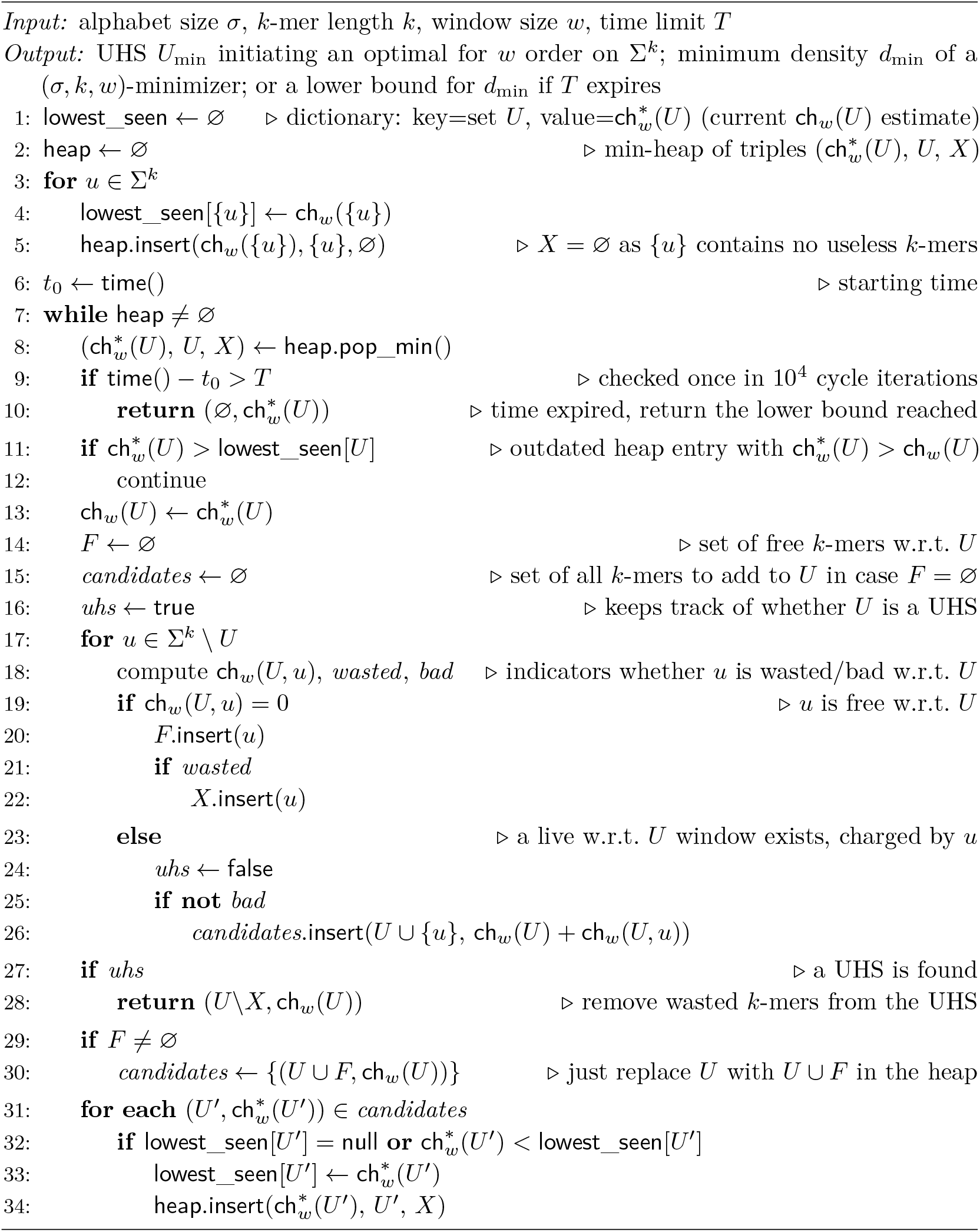

In our experiments, the heap operations seemed to have negligible effect on runtime. However, the space issue was critical. Two memory inefficiencies can be observed. First, since we insert triplets of the form 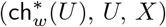 into the heap instead of updating 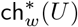 like in OM-basic /OM-phase, we store multiple entries with obsolete 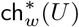 values (which are discarded in lines 11–12 of Algorithm 4). Second, the dictionary lowest _seen keeps accumulating entries during the course of the algorithm, even for sets that will never be used again. To cope with this increased memory usage we implemented two cleanup routines, called once every *M* iterations (we set *M* = 30,000 in our experiments). The first routine removes any heap element 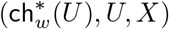 where 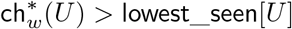. This is safe: the removed elements are never processed during the course of OM-heap. The second routine removes all lowest _seen entries with lower values than 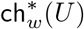 on the top of the heap. This is unsafe: if the entry by the key *U* is deleted, we later may reach *U* again (via a suboptimal arrangement). This will result in reinserting this key with a suboptimal value, and inserting *U* into the heap with the same suboptimal value. This increases the workload, leading to suboptimal performance, but cannot result in a wrong answer as the set *U* has been earlier processed with the correct value. In fact, the second cleanup is a tradeoff between the memory usage and the number of processed sets.

### A.3 ILP numerical results

GreedyMini minimizers (generated using default parameters) were optimal for many (*w, k*) pairs on which we ran ILP (Table A1). For instance, for *w* = 2 some GreedyMini orders were optimal for all values we tested up to *k* = 11 (the cases *k* ∈ {8, 10} did not terminate) and for *w* = 3 up to *k* = 4. The only pair of (*w, k*) for which an optimal minimizer could be generated by GreedyMini and then proven using ILP (without the KGMLT bound), but not generated by OM-phase /OM-heap within 10 hours, was (*w, k*) = (2, 6).

**Table A1.** Comparison of the number of charged windows for orders generated by GreedyMini, our ILP solver for minimizers, and Kille *et al*.’s ILP solver [8] for forward sampling schemes over *σ* = 2. All the results are optimal. Bold entries match the KGMLT lower bound.

| w | k | GreedyMini | ILP | FS optimal | #Windows |
| --- | --- | --- | --- | --- | --- |
| 2 | 2 | 11 | 11 | **10** | 16 |
| 2 | 3 | **20** | **20** | **20** | 32 |
| 2 | 4 | 38 | 38 | **37** | 64 |
| 2 | 5 | **74** | **74** | **74** | 128 |
| 2 | 6 | 144 | 144 | **143** | 256 |
| 2 | 7 | **286** | **286** | **286** | 512 |
| 2 | 9 | **1,118** | **1,118** | **1,118** | 2,048 |
| 2 | 11 | **4,412** | **4,412** | **4,412** | 8,192 |
| 3 | 2 | 16 | 16 | 15 | 32 |
| 3 | 3 | 29 | 29 | 29 | 64 |
| 3 | 4 | **56** | **56** | **56** | 128 |
| 3 | 7 | **411** | **411** | **411** | 1028 |
| 3 | 10 | **3,152** | **3,152** | **3,152** | 8,192 |
| 4 | 2 | 25 | 25 | 24 | 64 |
| 4 | 3 | 46 | 46 | 45 | 128 |
| 4 | 4 | 87 | 87 | 87 | 256 |
| 4 | 5 | **172** | **172** | **172** | 512 |
| 5 | 6 | **560** | **560** | **560** | 2,048 |

### A.4 Obtaining average and GreedyMini density results for Fig. 2

#### Average density

For *k* ≤ 4, we computed the average density exactly using Theorem 9. For *k* = 5, 6, we computed this density exactly for *w* ≤ 20, using [3, Theorem 1], and estimated it by a random sample of (*w*+*k*)-windows for *w >* 20, using [3, Lemma 1].

#### GreedyMini density

To complete the results for *k* = 6, we used the density of the best available minimizer, provided by the GreedyMini method [4], as the upper bound for the minimum density. For 2 ≤ *w* ≤ 15, we used the minimizers published in the GreedyMini repository^6^. In addition, we generated orders for up to *w* = 60 with the `GM-improve` tool of GreedyMini, iteratively using the order for *w* − 1 as the starting point and setting `-max_swapper_time_minutes` to 5.

## Footnotes

1 Such an assumption requires *σ*^*w*+*k*^ to fit into *O*(1) machine words. This condition holds in all cases feasible for OptMini, as *k* must be small and *w* = *O*(*σ*^*k*^) is a natural bound: for larger *w*, even the *average* density is close to the lower bound 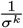 [3].

2 Since *U* is not necessarily a UHS **for** *w*, different such orders *ρ* can define different minimizers (*ρ, w*), but all of them have the same density.

3 At least one near-UHS is discovered as *C >* ch_*w*_ (Σ^*k*^).

4 In this case, the minimum density is known as the KGMLT lower bound is hit by a GreedyMini order.

5 By definition of wasted *k*-mer, the sets *U* and *U \ X* share the same set of live windows. Hence, if *u* is wasted w.r.t. *U*, then *u* is wasted w.r.t. *U \ X*.

6 https://github.com/OrensteinLab/GreedyMini

## Notes

**Funding** This study was supported by Israel Science Foundation grant no. 358/21 to Yaron Orenstein.

### Competing Interest Statement

The authors have declared no competing interest.

### Summary of Updates

Longer version to match current paper

